# Addressing the dynamic nature of reference data: a new nt database for robust metagenomic classification

**DOI:** 10.1101/2024.06.12.598617

**Authors:** Jose Manuel Martí, Car Reen Kok, James B. Thissen, Nisha J. Mulakken, Aram Avila-Herrera, Crystal J. Jaing, Jonathan E. Allen, Nicholas A. Be

## Abstract

**Background:** Accurate metagenomic classification relies on comprehensive, up-to-date, and validated reference databases. While the NCBI BLAST Nucleotide (nt) database, encompassing a vast collection of sequences from all domains of life, represents an invaluable resource, its massive size —currently exceeding 1012 nucleotides— and exponential growth pose significant challenges for researchers seeking to maintain current nt-based indices for metagenomic classification. Recognizing that no current nt-based indices exist for the widely used Centrifuge classifier, and the last public version was released in 2018, we addressed this critical gap by leveraging advanced high-performance computing resources.

**Results:** We present new Centrifuge-compatible nt databases, meticulously constructed using a novel pipeline incorporating different quality control measures, including reference decontamination and filtering. These measures demonstrably reduce spurious classifications, and through temporal comparisons, we reveal how this approach minimizes inconsistencies in taxonomic assignments stemming from asynchronous updates between public sequence and taxonomy databases. These discrepancies are particularly evident in taxa such as *Listeria monocytogenes* and *Naegleria fowleri*, where classification accuracy varied significantly across database versions.

**Conclusions:** These new databases, made available as pre-built Centrifuge indexes, respond to the need for an open, robust, nt-based pipeline for taxonomic classification in metagenomics. Applications such as environmental metagenomics, forensics, and clinical metagenomics, which require comprehensive taxonomic coverage, will benefit from this resource. Our new nt-based index highlights the importance of treating reference databases as dynamic entities, subject to ongoing quality control and validation akin to software development best practices. This dynamic update approach is crucial for ensuring the accuracy and reliability of metagenomic analysis, especially as databases continue to expand in size and complexity.

## Background

The capacity to accurately and sensitively profile microbial communities is critical to a broad swath of biological applications, spanning biomedical research, healthcare, and environmental biosurveillance. Microbial community features are indicative of a multitude of human health-relevant outcomes [1], including cancer [2], infection [3, 4, 5], trauma [6, 7], chronic injury [8, 9], and neurological disease [10, 11, 12]. Similarly, profiling microbial communities in non-human environments can provide detection and association information relevant to bioremediation, environmental surveillance [13, 14], pest control [15], and wastewater monitoring [16, 17]. In these applications spaces, the microbial community is often profiled via metagenomic sequencing, which facilitates comprehensive detection of fastidious or difficult-to-culture microorganisms. Metagenomic classification transforms raw shotgun DNA sequences into microbiome composition profiles, which can then be interrogated for signatures or biomarkers of any given label of interest.

A broad array of software platforms have been developed for classification of shotgun metagenomic sequence reads, including Kraken [18, 19], Centrifuge [20], MetaPhlAn [21], CLARK [22, 23], DIAMOND [24], Kaiju [25], GOTTCHA [26], metakallisto [27], KMCP [28], LMAT [29, 30], and ganon [31]. Each of these platforms relies, in some form, on a reference database containing known sequences or sequence signatures from a subset of reference genomes. These databases may include a range of indexed reference genomes, a subset of discriminative k-mers (e.g., CLARK and KrakenUniq [32]), or a curated set of marker genes (e.g., MetaPhlAn4 [21]). The selection of database and software platform has a clear and systematic impact on the resultant microbial profile, influencing downstream assessments [33]. Numerous additional benchmarking studies providing comparative assessments between platforms are available for assessment [34]. In addition to the reference source and taxonomic clades for inclusion, metagenomic classification results are impacted by temporal distribution of build dates and filtering processes applied to the reference content. These parameters are potentially impactful but have not been thoroughly assessed in previous studies.

To address this knowledge gap, we undertook a comparison of five temporally distinct database builds, comparing the extent of read classification and overall compositional impact. We also assessed the impact of reference decontamination via Conterminator [35] and Recentrifuge [36] platforms and the impact of short reference sequence removal. For broad taxonomic coverage, we constructed databases from the NCBI BLAST Nucleotide (nt) database [37], the most comprehensive database for nucleotide BLAST search [38]. We selected the Centrifuge classifier platform for this comparison due to its classification speed and optimized indexing scheme.

The NCBI BLAST nt database encompasses nearly all traditional GenBank divisions, representing a significant portion of available GenBank sequences and spanning all domains of life [37, 39]. Therefore, the growth rate of nt follows that of GenBank. Both the number of sequences and the nucleotides in GenBank are experiencing an exponential growth (see Figure 2 for quantitative details). Building nt-based reference databases for taxonomic annotation requires increasing computational resources, such as processing power, memory, and I/O bandwidth. The build process for these databases is particularly memory-intensive, requiring specialized computing architectures often not readily available to most researchers. This has led to increasing reports of difficulties and failures in building such databases as dataset sizes continue to grow [31, 42]. Prior to this effort, the latest available Centrifuge database based on NCBI BLAST nt was built in 2018, with multiple users unsuccessfully requesting an updated version in recent years [43]. As a response to the need for an open, robust, nt-based pipeline for taxonomic classification in metagenomics, we provide updated, stable, and optimized Centrifuge nt reference databases.

**Figure 1.**
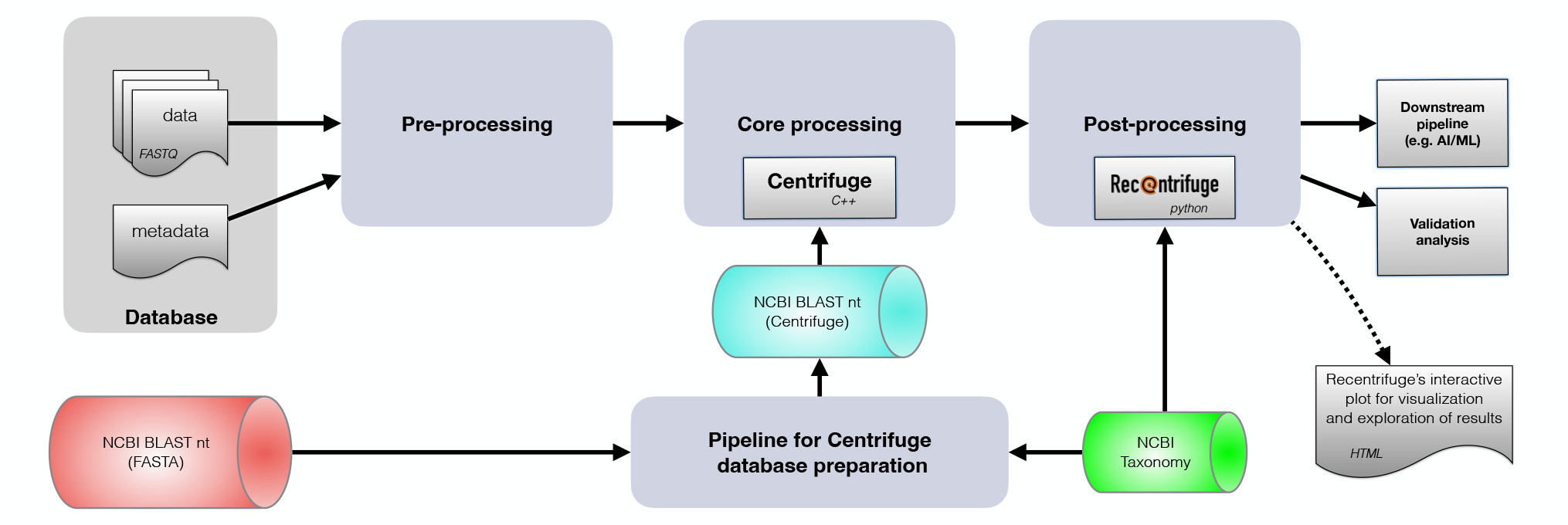
General layout of a metagenomic classification pipeline based on an NCBI BLAST nt database using Centrifuge and Recentrifuge. This flowchart is showing a typical configuration of a pipeline for taxonomic classification in metagenomics using the NCBI BLAST nt database [37, 39]. Data (and metadata, as available) is feed into a FASTQ format pre-processing block that performs quality checks using software such as fastp [40] and MultiQC [41]. Filtered FASTQ files are then processed by Centrifuge [20] using an indexed version of the nt database. Finally, Recentrifuge [36] post-process Centrifuge output, including negative control samples (if any), to further filter and prepare results for interactive exploration, downstream analysis and validation tests. Typical downstream jobs include metagenome assemblers and artificial intelligence/machine learning (AI/ML) training pipelines. The pipeline for building the Centrifuge reference databases based on nt is detailed in Figure 3.

**Figure 2.**
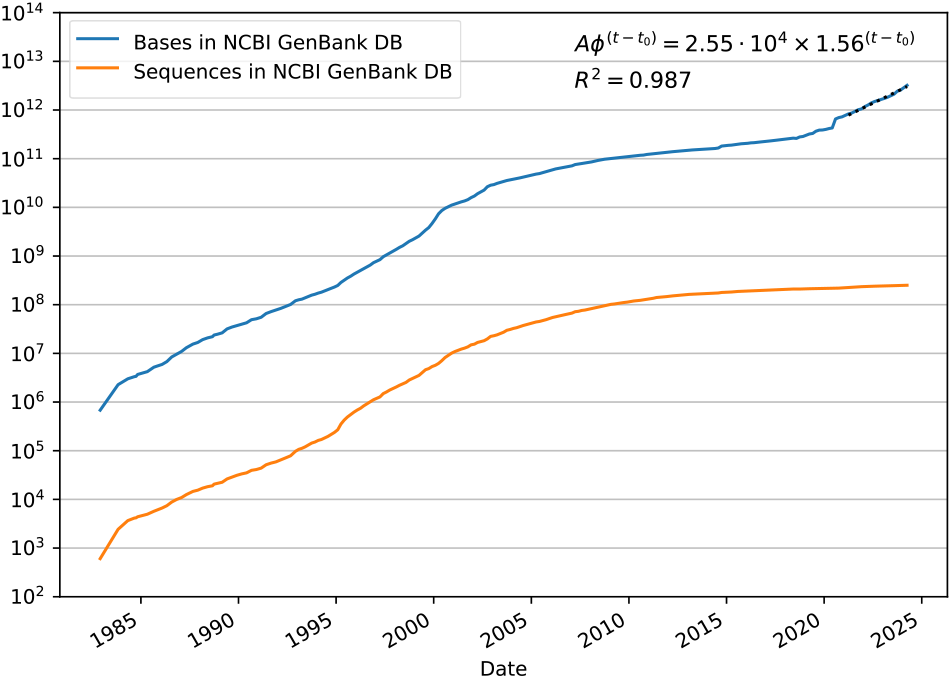
Evolution of NCBI GenBank database over time. This semi-log plot shows the number of bases and sequences for NCBI GenBank. For the last 3 years of the time series, from April 2021 to April 2024, we fit an exponential model 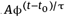 (with time constant τ = 1) to the number of bases in the database, with *R*^2^ confirming the goodness of the model. As time *t* (x-axis, date) is measured in years from *t*_0_ = 1982, *A* is the intercept to *t* = *t*_0_ and ϕ is the base of the exponential with the meaning of the yearly rate of change [44]. So, on average, between the mentioned dates, GenBank database had a 56% growing rate in nucleotides from year to year. In addition, at least in the last decade, the slope of the number of bases has been increasing while the slope of sequences has been decreasing, with these opposite trends indicating a progressive rise in the average length of the sequences in the database.

The updated databases provide a critical resource for researchers using metagenomics, where comprehensive and reliable taxonomic classification across the tree of life is paramount. This work underscores the critical need for treating these databases as a dynamic entity requiring continuous quality control and validation, much like software development best practices.

## Methods

### Reference selection

Five NCBI nt release dates spanning 2022-2023 were selected for assessment (Table 1).

**Table 1.** Size parameters for nt DB references and builds.

| nt DB release date | Build version name | Masked nt DB size |  | Centrifuge nt indexed DB size |  |  | Taxonomy DB |  |
| --- | --- | --- | --- | --- | --- | --- | --- | --- |
|  |  | std | dec | std | dec | filt | size | content |
| Jun 26, 2022 | Jun22 | - | 745 GiB | - | 442.02 GiB | - | 203 MiB | 2.43 Mtid |
| Sep 23, 2022 | Sep22 | - | 843 GiB | - | 498.89 GiB | - | 213 MiB | 2.45 Mtid |
| Jan 04, 2023 | Jan23 | 941 GiB | 936 GiB | 556.77 GiB | 553.66 GiB | 570.34 GiB | 217 MiB | 2.48 Mtid |
| Apr 03, 2023 | Apr23 | 1.09 TiB | 1.08 TiB | 614.15 GiB | 610.53 GiB | - | 220 MiB | 2.50 Mtid |
| Oct 17, 2023 | Oct23 | - | 1.30 TiB | - | - | 784.39 GiB | 227 MiB | 2.53 Mtid |

In the database size columns, **std** refers to a standard database (non-decontaminated), **dec** denotes a decontaminated database while **filt** is a decontaminated database that, in addition, has been processed by Recentrifuge's *refasfilt* script.
The size of taxonomy database is the decompressed size in megabytes of the *names.dmp* file of the corresponding NCBI Taxonomy database available at the indicated nt database release date.
The unit tid denotes (different) taxonomic identifiers, so Mtid is a million of taxonomic identifiers.

### Characterization and decontamination of the reference database

Figure 3 shows a flowchart of the main steps involved in the pipeline for building a reference database for Centrifuge based on the nt database.

**Figure 3.**
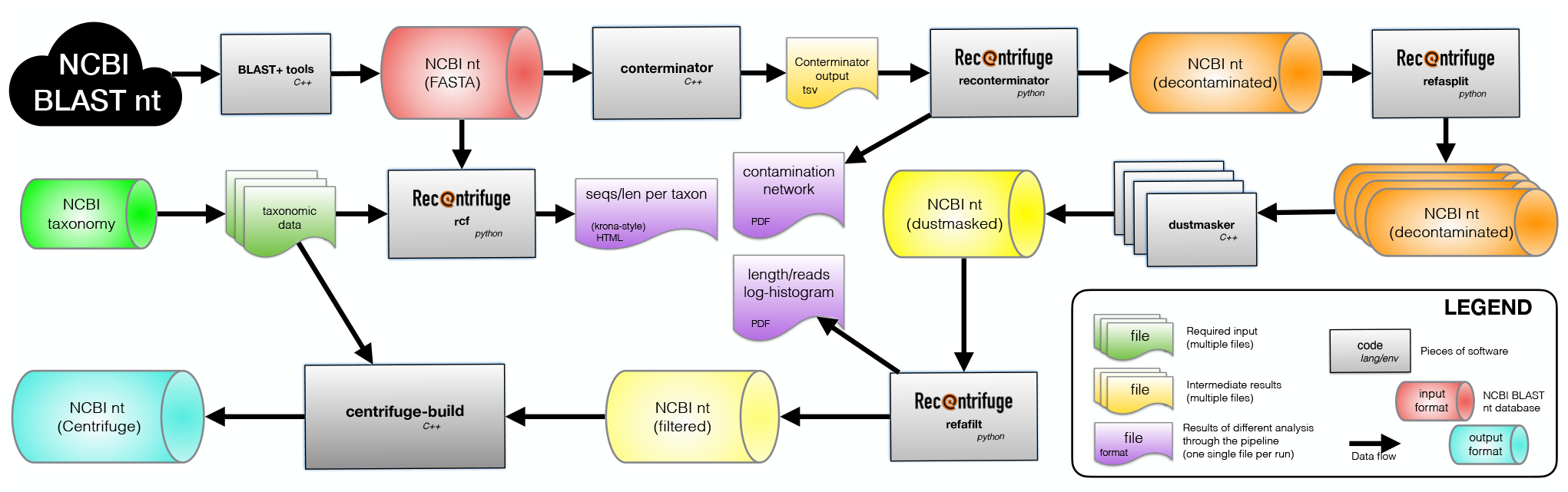
General flowchart of the pipeline for building the Centrifuge reference database based on the NCBI BLAST nt database. The entire pipeline process takes several weeks using a HPC (high-performance computing) server with a high-memory-per-core ratio, a high-bandwidth low-latency parallel filesystem, 128 cores in two AMD EPYC 7742 64-core processors at 2250 MHz base frequency, with 512 MiB of cache, and total memory of 2 TiB. The most demanding step is the final indexing of the filtered database using a version of centrifuge-build, which may expand beyond two weeks, followed by the decontamination step, which usually takes a few days.

After obtaining the nt database in FASTA format using BLAST+ tools [45] and the current content of the NCBI Taxonomy database, the content of nt was explored using Recentrifuge [36]. Figures 4 and S1 show some examples of insights into the database provided at this step.

**Figure 4.**
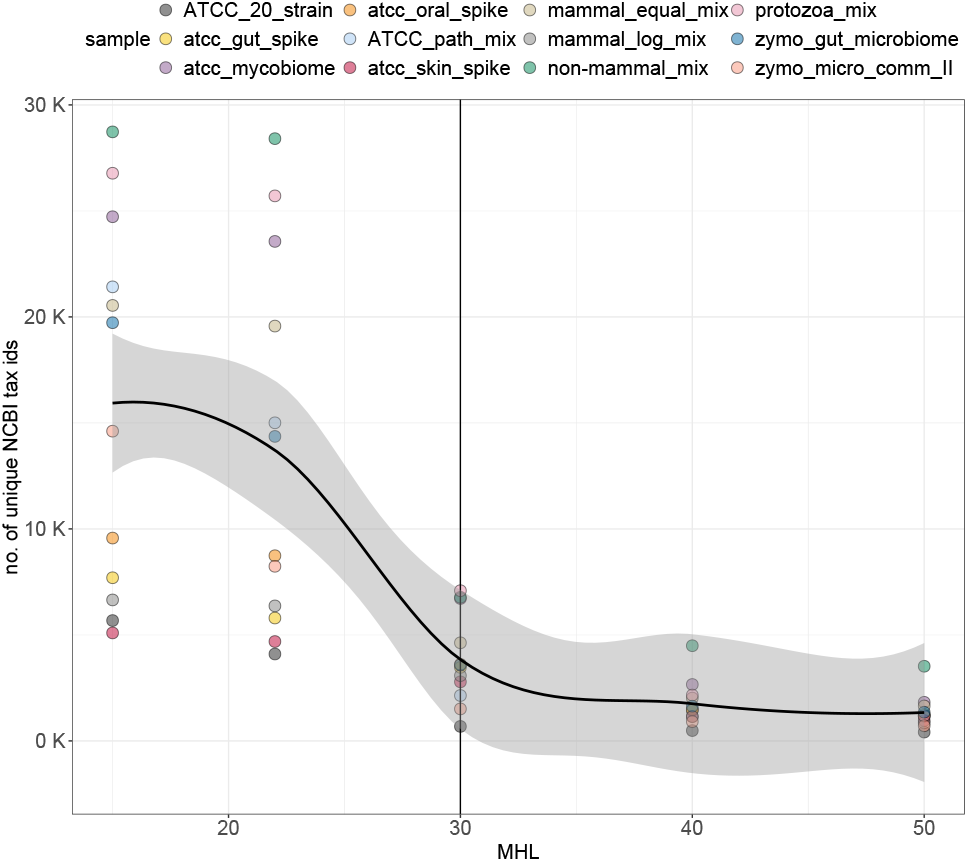
The number of unique NCBI tid captured at different MHL values across samples and averaged across all temporal database versions. Data points are colored according to sample.

The nt database was decontaminated using Conterminator [35] in parallel and the output post-processed by a different script of Recentrifuge, which removes the detected contamination and generated a contamination network plot (see Figure S2 for an example).

### Reference masking, filtering, and indexing

The `refasplit` script from Recentrifuge [36] was used throughout the pipeline to facilitate efficient parallel I/O processing of the sequence database, which contains on the order of 1012 nucleotides. Low-complexity regions were masked using several instances of NCBI dustmasker [45] in parallel. The decontaminated and masked version of the nt database was then processed by Recentrifuge’s `refafilt` code to remove sequences shorter than 16 nucleotides, which were found to cause downstream issues, and to demultiplex headers for redundant sequences as generated by recent versions of the BLAST+ command line tool `blastdbcmd`. Finally, indexing of the decontaminated filtered database was performed with an optimized version of the Centrifuge 1.0.4-β database building C++ code `centrifuge-build` [20] targeting Livermore Computing (LC) HPC cluster nodes with 128 cores and 2 TiB main memory.

The entire build process, which demanded substantial computational resources, required more than three weeks of clock time. This is equivalent to roughly 6 years of continuous computation on a single processor, underscoring the immense computational effort involved.

### Sample sequencing and processing

Defined DNA mixtures were employed for comparative assessment of databases and classification parameters (Tables 1, 2). No-template process negative controls (ntc) were included in all analyses. Sequence libraries were prepared from gDNA via the Illumina DNA Prep library preparation kit and sequenced on the Illumina NextSeq 2000 via P2 300 cycle reagents and flow cell. Resultant data were processed for taxonomic classification via Centrifuge using each of the database builds described in Table 1. Comparative analyses of reference controls were carried out in R (version 4.2.0). Phyloseq [46] was used to generate microbiome abundance profiles from count data and ggplot2 [47] was used to generate figures. The vegan package [48] was used to compute Bray Curtis dissimilarity distances between samples.

**Table 2.** DNA standard mixtures employed for comparative assessment.

| Sample name | Mixture | Vendor | Cat No | Human spike | Sample type |
| --- | --- | --- | --- | --- | --- |
| zymo_gut_microbiome | Zymobiomics gut microbiome | Zymo | D6331 | no | bacteria |
| zymo_micro_comm_II | Zymobiomics microbial community standard II | Zymo | D6311 | no | bacteria |
| atcc_mycobiome | Mycobiome mock community | ATCC | MSA-1010 | no | eukaryote |
| atcc_oral_spike | Oral microbiome mock community | ATCC | MSA-1004 | 18 ng | bacteria |
| atcc_skin_spike | Skin microbiome mock community | ATCC | MSA-1005 | 14 ng | bacteria |
| atcc_gut_spike | Gut microbiome mock community | ATCC | MSA-1006 | 40 ng | bacteria |
| protozoa_mix | *Acanthamoeba castellanii*, *Naegleria fowleri* | ATCC | Various | no | eukaryote |
| mammal_equal_mix | Equal mix of 11 mammalian DNAs | Various | Various | no | eukaryote |
| non-mammal_mix | Equal mix of 10 non-mammalian eukaryotic DNAs | Various | Various | no | eukaryote |
| mammal_log_mix | Log concentration range of 7 mammalian DNAs | Various | Various | no | eukaryote |
| atcc_20_strain | ATCC 20 strain mix | ATCC | MSA-1003 | no | bacteria |
| atcc_path_mix | ATCC pathogen mix | ATCC | MSA-4000 | no | bacteria |

## Results and Discussion

### Comparative assessment of gDNA reference controls and mixtures

In addition to the assessment of temporal database builds, we evaluated other microbiome analytical parameters relevant to the assessment of accurate microbiome profiles. This warrants the use of standard controls with known microbial composition across different kingdoms of life (Tables 1, 2). We systematically evaluated the impact of different values for the minimum length of partial hits in Centrifuge [20] or MHL (Minimum Hit Length), relative abundance thresholds, taxonomic levels, and sample type to generate accurate microbiome profiles. We calculated precision, recall, balanced F1 scores and Bray Curtis dissimilarity distances to compare sequenced profiles of control standards with ground truth data across all database versions (Table 2).

First, we investigated the impact of temporal database builds on microbiome profiling and observed temporal instability in Bray Curtis dissimilarity distances in 3 samples; ATCC_path_mix, proto-zoa_mix and zymo_micro_comm_II when compared to reference control profiles (Figures 4, 5, and S3). There was an increase in Bray Curtis dissimilarity distances in the Jan23 database versions for these samples. Upon inspection, we determined that delayed updates between taxonomic labels in NCBI BLAST nt database and the current NCBI Taxonomy database at the time of the nt database release was a major factor. The nt database is relying on the Taxonomy database for the taxonomic information, but both databases are evolving over time independently, with no explicitly version mapping between them in the documentation. Traditionally, the nt database was using a current version of the taxonomy database but that does not seem the case anymore, with the taxonomy used in NCBI BLAST nt database slightly trailing behind. In some cases, newly uploaded genomic sequences were lacking current taxonomic labels and then confounded Centrifuge’s database indexing algorithm and, eventually, its classification process.

**Figure 5.**
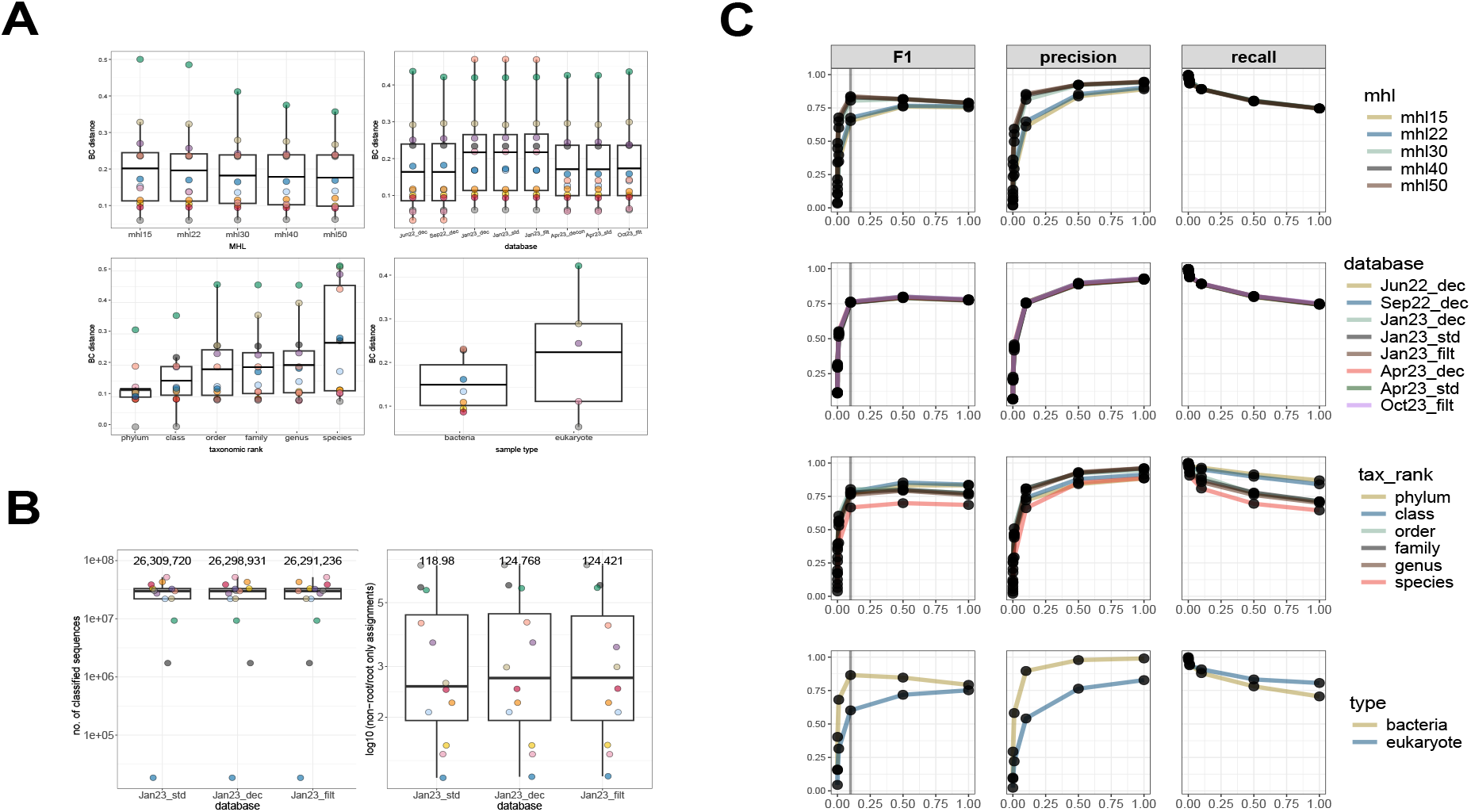
Comparative assessment of gDNA reference controls. A) Comparison of Bray Curtis dissimilarity distances at an abundance threshold of 0.1% between MHL values, database versions, taxonomic ranks, and sample types. For every variable that is being compared, Bray Curits dissimilarity distances were averaged across other variables. B) The number of annotated sequences and ratio of non-root to root-only assigned reads across different versions of the Jan23 database. Mean values are annotated at the top of each boxplot. C) Comparison of classification accuracy metrics (F1 balanced accuracy, precision, and recall) between MHL values, database versions, taxonomic ranks, and sample types across abundance thresholds.

For example, in the protozoa_mix sample, there were reads that were assigned and equally scored to multiple *Naegleria fowleri* (tid 5763) reference genomes that were present in the Jan23 NCBI BLAST nt database. However, when Centrifuge was reporting multiple assignments (Centrifuge default), only reads that were assigned to *N. fowleri* genomes whose tid was present in the NCBI Taxonomy database used during the database compilation were assigned to tid 5763 while reads that were assigned to *N. fowleri* genomes whose tid was absent from the used taxonomy (but present in the NCBI BLAST nt database) were classified with a tid 0, normally reserved for unclassified reads. When Centrifuge was configured to report a unique tid per read using the lowest-common-ancestor (LCA) strategy (pipeline default), those reads were given a final root-only assignment (tid 1). As another example, the results for the zymo_micro_comm_II sample were impacted when using the Apr23 and Oct23 database due to varied taxonomic classification of *Listeria monocytogenes* genomes (Figures 5A, S3). In that database, the addition of a *Listeriamonocytogenes* genome (CP046449.1) annotated as Listeria sp. LM90SB2 (tid 2678528 in the NCBI Taxonomy database, with parent being unclassified *Listeria*, tid 2642072) resulted in a final LCA classification of reads at only the genus level (*Listeria*). Overall, our observations highlight the impact on the classification of specific taxa due to the discrepancy of taxonomic information between the nt database and the version of the NCBI Taxonomy database used for indexing the Centrifuge database.

Next, we tested the impact of the following MHL values: 15 (minimum value), 22 (Centrifuge’s default), 30, 40, and 50 on the number of unique NCBI tax ids assigned to sequenced reads. Here, we observed that the number of unique NCBI tax id assignments and Bray Curtis dissimilarity distances decreases at increasing MHL values (Figures 4 and 5A). An inflection point was observed at MHL 30 (Figure 4), followed by an increase in F1 and precision scores compared to MHL 15 and 22 (Figure 5C). This indicates that a MHL of 30 is likely close to the optimal value in reducing false positive assignments across these particular samples (Figure S3A) by reducing the noise of spurious assignments without a significant increase in false negatives.

As most microbiome analysis protocols remove low-abundance features to account for potential sequencing errors and false positives, we assessed the impact of different thresholds (0, 0.001, 0.01, 0.1, 0.5, 1) on classification performance. Overall, we observed that a relative abundance threshold of 0.1% resulted in the highest F1 scores across samples (Figure 5C). As expected and previously observed, our data demonstrated that classification performance decreases at lower taxonomic ranks with an increasing number of detected false positives (Fig. S3A). Furthermore, samples consisting mainly of bacterial content had a better classification performance than eukaryotic samples (Figure 5B,C).

Finally, we compared the number of annotated sequences between the different versions of the Jan23 database and observed a decrease along the implemented improvements (Fig 5B). The standard version (Jan23_std) had an average number of 26,309,720 annotated sequences, followed by the decontaminated version (Jan23_dec) with 26,298,931 annotated sequences and lastly, the decontaminated and filtered version (Jan23_filt) had 26,219,236 annotated sequences. However, Jan23_dec and Jan23_filt databases had higher non-root to root-only assignment ratios, suggesting that additional annotated reads with the standard version (Jan23_std) were assigned to the root-only level.

## Potential implications

While this work primarily focuses on providing a readily usable and improved nt-based database for metagenomic classification with Centrifuge and Recentrifuge, the underlying methodology and findings have implications that extend beyond this immediate application.

Firstly, the availability of a comprehensive, decontaminated nt database opens new avenues for developing and training machine learning models for metagenomic analysis. Such models, trained on cleaner data, have the potential to achieve higher accuracy and better generalization capabilities, particularly for challenging tasks such as identifying novel or rare taxa. This could lead to improved tools for metagenome-based diagnostics, discovery of novel microbial enzymes, and understanding the complex interactions within microbial communities.

Secondly, our observation of temporal inconsistencies in taxonomic assignments stemming from asynchronous updates between NCBI databases highlights a crucial challenge in metagenomic analysis. It underscores the need for a paradigm shift in how we approach reference data, moving from static resources to dynamically maintained and versioned collections. Drawing inspiration from software development best practices, continuous integration and continuous delivery (CI/CD) pipelines could be adapted for reference databases, ensuring timely updates, robust validation, and improved reproducibility across studies.

Additionally, the pipeline developed for database construction, encompassing decontamination, filtering, and validation steps, can be generalized to other large reference databases commonly employed in metagenomic analysis. Applying similar quality control measures to resources like the NCBI nr (non-redundant protein) database or specialized databases for viral or fungal identification could significantly enhance the accuracy and reliability of classification in those domains. This has the potential to improve research outcomes in fields as diverse as environmental monitoring, disease diagnostics, and evolutionary biology.

Furthermore, the availability of a comprehensive, up-to-date, and validated database encompassing all domains of life—such as the one presented here—offers a unique advantage for researchers in fields that require analysis of complex microbial communities across diverse environments. For example, studies investigating host-microbe interactions, exploring the microbiome of underexplored ecosystems, or tracing the origins of emerging pathogens can benefit greatly from such a resource.

However, the exponential growth of sequencing data and the associated computational burden highlight the need for innovative strategies to ensure the long-term sustainability of such comprehensive resources. This might involve exploring distributed computing approaches, developing more efficient indexing and search algorithms, splitting the database, or even dividing the classification problem, e.g., converting the classification goal in a hierarchical problem by using a “forest” of specific classifiers to decide the annotation at different taxonomic levels instead of a single classifier covering the entire tree of life.

## Data availability

As of September 2024, we have released two decontaminated and filtered Centrifuge nt databases: Jan23_filt (`nt_wntr23_filt`) and Oct23_filt (`nt_fall23_filt`). The databases can be freely downloaded from AWS storage following the instructions at Langmead Lab Centrifuge indexes webpage. Alternatively, they can be downloaded from LLNL Green Data Oasis (GDO) public ftp server at ftp://gdo-bioinformatics.ucllnl.org/centrifuge/nt_wntr23 using any software supporting anonymous ftp download. To ease the download process, the databases are split in ultra-compressed `7z` files of 4 GiB or less with name format `nt_wntr23_filt.cf.7z.*`. There are 71 `7z` files for the Jan23_filt database, and 99 `7z` files for the Oct23_filt version of the database. The `curl` command may be used to easily download those files and the `7z` command to extract the Centrifuge database files.

The corresponding taxonomy files for the databases, useful for Recentrifuge post-processing of Centrifuge results, are available at ftp://gdo-bioinformatics.ucllnl.org/centrifuge/nt_wntr23/ taxonomy for the Jan23_filt version of the database (substitute `nt_wntr23` with `nt_fall23` in the link to access the corresponding files for the Oct23_filt version). The following command may be used to download the taxonomy files:

~~~
curl “{full_taxonomy_url}/{names,nodes}.dmp” -O
~~~

where `{full_taxonomy_url}` should be substituted by the full ftp url just indicated for the taxonomy files.

While feasible, we plan to publicly release newer versions of the nt-based Centrifuge index we present in this article.

Shotgun metagenomic sequences of standard controls are uploaded to NCBI SRA with the Bioproject ID PRJNA1118346.

### Practical notes

Although the most demanding step is the indexing of the nt database as described above, using these large indexes with Centrifuge has important computational requirements, especially regarding free disk space and available main memory. When Centrifuge loads the Jan23_filt (`nt_wntr23_filt`) database into main memory, it can use close to 600 GiB. Given that size, a computer with at least 768 GiB of main memory is recommended for running the Centrifuge classifier. When using a high number of parallel threads (such as 256), 1 TiB of main memory is advisable. For the Oct23_filt (`nt_fall23_filt`) database, which is roughly 40% larger than Jan23_filt (1.4 trillion vs 1.0 trillion nucleotides in the source database), using a server with more than 1 TiB of main memory is unavoidable.

In addition, depending on the speed of the storage and memory access, Centrifuge may take more than half an hour just for loading the nt database from disk into memory. Given this overhead, it is recommended to use a sample sheet file (`–sample-sheet` argument in Centrifuge) to process multiple samples without requiring additional database loads.

Also, when replicating our pipeline, it is strongly recommended to use the LCA strategy with Centrifuge, by using the `-k 1` argument in the call to the classifier.

Finally, it is straightforward to use Recentrifuge to post-process Centrifuge’s results, with detailed documentation in Recentrifuge’s wiki at https://github.com/khyox/recentrifuge/wiki/Running-recentrifuge-for-Centrifuge.

## Declarations

## List of abbreviations

DB: Data Base
GDO: Green Data Oasis
HPC: High Performance Computing
I/O: Input/Output
LCA: Lowest Common Ancestor
MHL: Minimum Hit Length
WGS: Whole-Genome Shotgun

## Acknowledgements

We gratefully acknowledge Benjamin Langmead (Johns Hopkins University) for hosting our Centrifuge database on the AWS Public Dataset Program space, making this resource readily accessible to the research community in the Centrifuge indexes webpage. We also extend our thanks to Li Song (Dartmouth College) and Daehwan Kim (University of Texas), authors of the Centrifuge software, for their valuable contributions to this project.

## Supplementary figures

**Figure S1.**
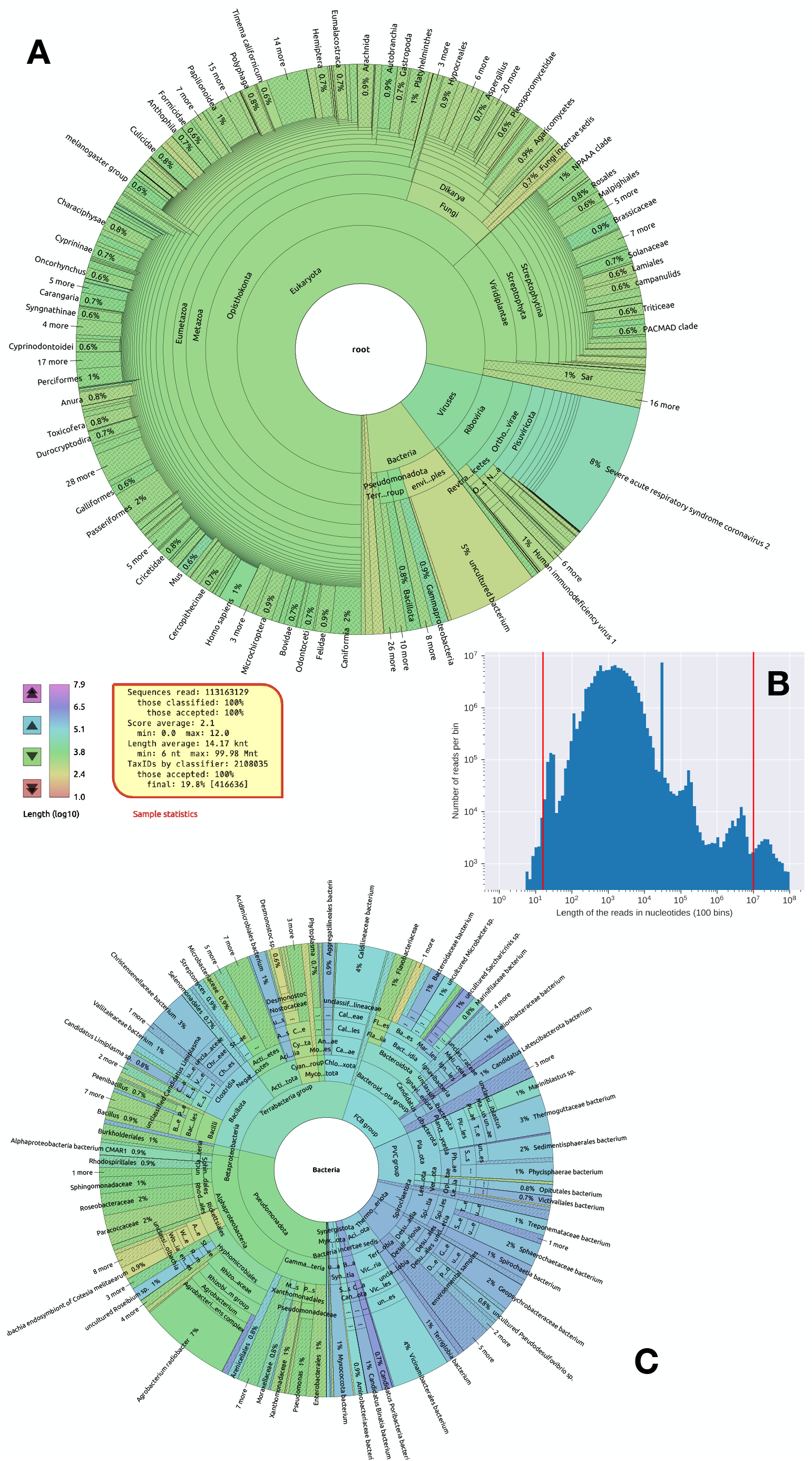
Recentrifuge’s scripts allow the characterization and interactive exploration of the content of the nt database during the building process of the nt database for Centrifuge. (A) Recentrifuge’s `rcf` interactive plot showing the content of the nt database for the number of sequences and the logarithm of their length in the color scale. (B) Recentrifuge’s `refafilt` output showing the distribution of the sequence lengths in the nt database. (C) Recentrifuge’s `rcf` interactive plot displaying new bacterial taxa not present in the previous seasonally compiled database showing the number of sequences and the logarithm of their length in the color scale.

**Figure S2.**
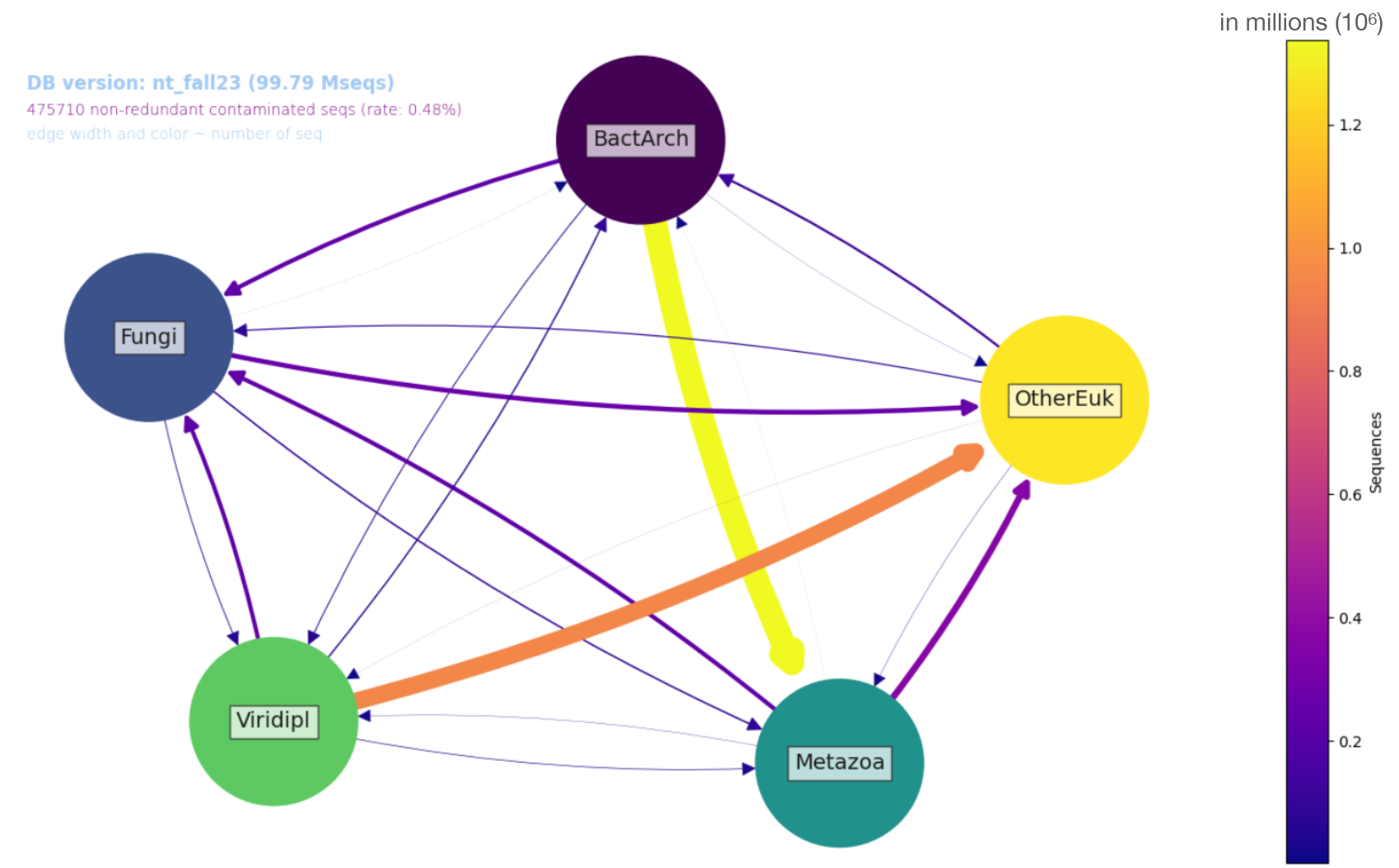
Example of inter-kingdom contamination network plot generated during the process of building the nt database for Centrifuge. The contamination was detected between the five default “kingdoms” as defined in Conterminator software, with the width and color scale of the arrows related to the number of sequences involved. The rate of contamination detected in the database (non-redundant sequences) before contamination removal was significant: almost half a percentage point.

**Figure S3.**
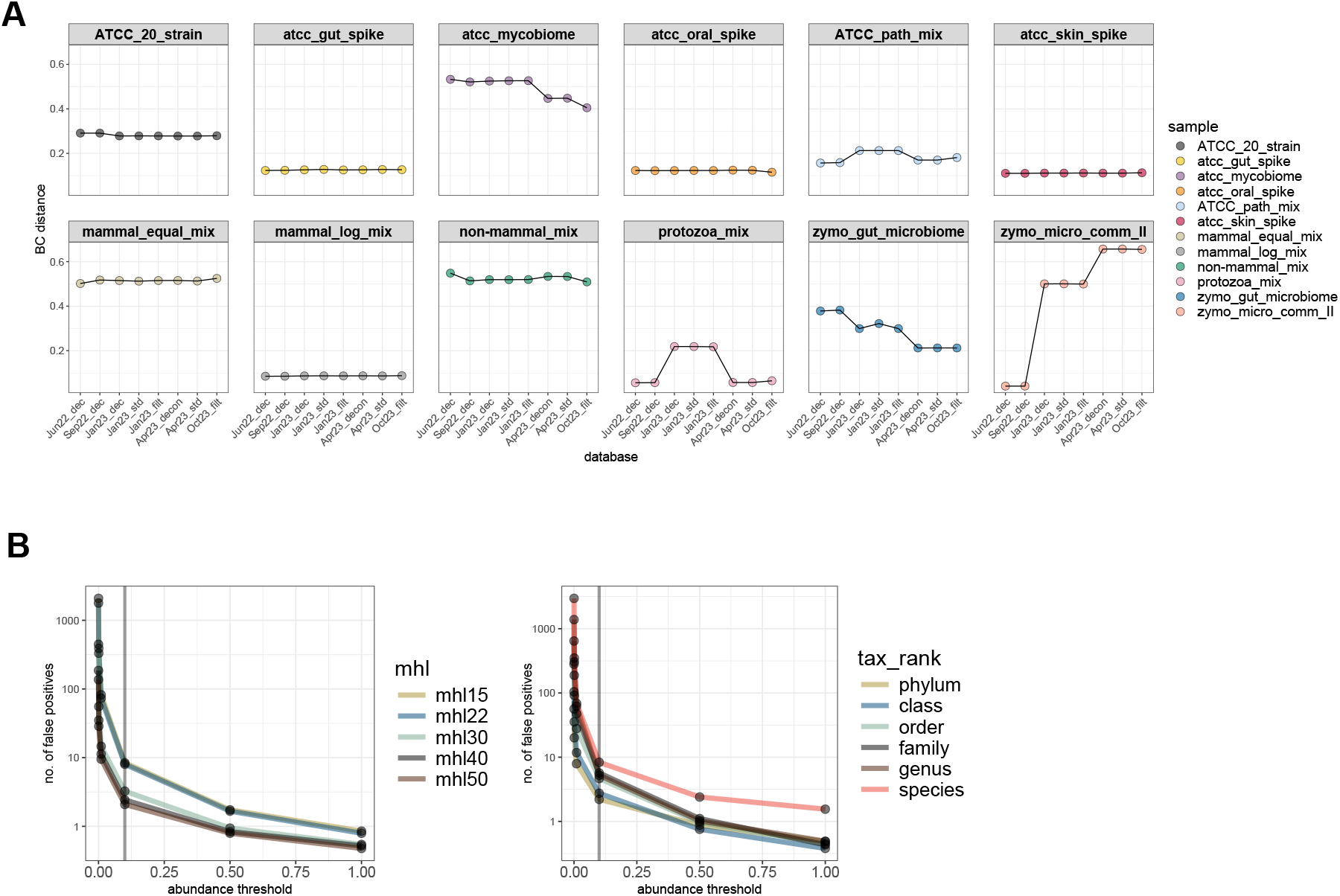
Additional comparative assessment of gDNA reference controls. A) Species-level Bray Curtis dissimilarity distance between database versions across all gDNA reference control samples. B) Number of false positive assignments between MHL values and taxonomic rank across abundance thresholds.

